# Characterizing a New Fluorescent Protein for Low Limit of Detection Sensing in the Cell-Free System

**DOI:** 10.1101/2022.04.06.487419

**Authors:** Caroline E. Copeland, Jeehye Kim, Pearce L. Copeland, Chloe J. Heitmeier, Yong-Chan Kwon

## Abstract

Cell-free protein synthesis-based biosensors have been developed as highly accurate, low-cost biosensors. However, since most biomarkers exist at low concentrations in various types of biopsies, the biosensor’s dynamic range must be increased in the system to achieve the low limits of detection necessary while deciphering from higher background signals. Many attempts to increase the dynamic range have relied on amplifying the input signal from the analyte, which can lead to complications of false positives. In this study, we aimed to increase the protein synthesis capability of the cell-free protein synthesis system and the output signal of the reporter protein to achieve a lower limit of detection. We utilized a new fluorescent protein - mNeonGreen, which produces a higher output than those commonly used in cell-free biosensors. Optimizations of DNA sequence and the subsequent cell-free protein synthesis reaction conditions allowed characterizing protein expression variability by given DNA template types, reaction environment, and storage additives that cause the greatest time constraint on designing the cell-free biosensor. Finally, we characterized the fluorescence kinetics of mNeonGreen compared to the commonly used reporter protein, superfolder Green Fluorescent Protein. We expect that this finely tuned cell-free protein synthesis platform with the new reporter protein can be used with sophisticated synthetic gene circuitry networks to increase the dynamic range of a cell-free biosensor to reach lower detection limits and reduce false positives proportion.

## INTRODUCTION

The cell-free protein synthesis (CFPS) system has been proven as a powerful platform for advancing our ability to study, exploit, and expand the potential of applied biotechnology and synthetic biology ^*1, 2*^. With the system’s unprecedented level of freedom and modularity to modify and control biological systems, the CFPS system allows for the prototyping of complex cellular functions by breadboarding genetic parts ^*3-9*^, genetic circuits ^*10-18*^, protein modification ^*19-22*^, and biosynthetic pathway ^*23-26*^. These advantages unlock the opportunity to transform the system into a versatile in vitro biosensing platform. Sensing biological artifacts such as nucleic acids ^*27-29*^, hormones ^*30*^, vitamin levels ^*31*^, harmful chemicals and compound levels ^*32-35*^, and heavy metals ^*35, 36*^, protein biomarkers ^*37, 38*^ and protein-protein interactions ^*39*^ can be done precisely, quickly, and inexpensively when utilizing the cell-free system (CFS).

These cell-free biosensors share the same core component of RNA and protein synthesis (transcription and translation), but differ in the way they detect the analyte and the cascade of events that happen between the detection, and RNA and protein synthesis. Cascade triggering methods and their targets include transcription factors to detect harmful small molecules, hormone receptors to detect endocrine disruptors, antibody-DNA conjugations to detect biomarker proteins, CRISPR-Cas proteins to differentiate between species variants, and riboregulators like the toehold switch to detect RNA associated with illnesses or riboregulators to detect fluoride ^*40*^. Researchers often use combinatorial methods in the cascade, with the entire sequence of events being classified as a gene circuit.

A low limit of detection (LOD) is a crucial feature for developing CFS biosensors because biomarkers and other molecules of interest are often present at very low levels in various types of specimens ^*41*^. Due to the frequent false-positive signals of low-cost, on-demand biosensors, researchers are required to add costly, time-consuming sample preparation processes. Another problem biosensors can come across is maximizing the dynamic range, which is the ratio between the leaky expression and the maximum expression (signal-to-noise ratio). Having a larger dynamic range will help the biosensor reach a low LOD and allow the sensor to pick up on a larger range of low levels of analyte. For transcription factor (TF)-based cell-free biosensors, decreasing the amount of TF or increasing the amount of reporter DNA with the operator can aid in reaching a low LOD, but then the dynamic range could be lost ^*34, 35*^. Others that have had problems with dynamic range have had to lower their reporter DNA concentration, which lowers the LOD ^*37*^. To increase dynamic range, methods have been created to scale up the cascade input from the target molecule to a higher detectable level, known as genetic amplifiers or positive feedback loops. Examples of amplifiers include the activation of robust ligand-free TFs ^*42, 43*^, a TF-free bacteriophage RNA polymerase or sigma factor-endogenous RNA polymerase pair ^*44*^, an anti-repressor RNA aptamer ^*35*^, or by a tag-specific protease targeting a repressor ^*45*^. The issue with adding an amplifier feedback loop or an amplification step of the analyte is that if either one is triggered falsely, the signal will be much higher than what it would have been without that extra step.

One type of amplifier with the potential for a false positive signal includes nucleic acid-based amplifiers, especially those that are isothermal reactions. These are very common for nucleic acid sensors because the DNA/RNA of interest can trigger a PCR-like reaction. One popular example of a sensor that uses an amplifier includes the toehold switch that detects Zika virus from serum and uses nucleic acid sequence-based amplification (NASBA) as an amplification step to reach a low enough LOD to detect the virus concentration in human serum at 7.2 × 10^5^ copies/ml (1.2 fM) ^*46*^. However, NASBA can create off-target amplification from a human serum sample full of other RNA molecules because of the difficulty of efficient primer binding to RNA ^*47*^.

If off-target amplification occurs, amplicons of any length can be made, especially since this is an isothermal system, taking away the barrier of the extension time, allowing extension to occur for the duration (3 hours), as well as the annealing temperature sequence specificity barrier. The toehold switch can be triggered by a sequence of 11% similarity to the target binding region (4-nt). Experimentally, they demonstrate low dynamic range when detecting Zika virus in macaque serum, since the background signal from heavily purified healthy serum, which was purchased from a manufacturer, had 30% expression compared to the reaction with infected macaque serum and signaling nearly at the same time, with probably lower dynamic range if healthy macaque serum was used as a control instead ^*46*^.

Here we aimed to amplify the output signal in other ways that do not involve the analyte, but rather by increasing the signal of the output protein and maximizing the performance of the cell-free protein synthesis reaction overall. To achieve the overarching goal of this study, we investigated various cell-free conditions and components that can potentially improve cell-free biosensor development. One of the largest contributions to cell-free biosensors in this research involves introducing the robust fluorescent protein, mNeonGreen (mNG), which has been highlighted as the brightest fluorescent protein ^*48, 49*^. We achieved a 2.6 times higher signal from mNG than the commonly used superfold green fluorescent protein (sfGFP) in the CFS. We also compared maturation time and fluorescence output rate to evaluate if mNG is comparable to the sfGFP ^*50*^. We investigated the DNA templates by optimizing different sequence elements and characterizing protein expression by template types. We assessed the ribosome binding site, 5’- untranslational region (UTR), spacer sequence, and codon usage to measure the DNA template-dependent cell-free protein synthesis capacity. In addition, we evaluated the cell-free protein synthetic tolerance on various additives and environmental matrix effects. We anticipate that the finely tuned CFS platform in this study can be used with sophisticated synthetic gene circuitry networks to increase the dynamic range of a cell-free biosensor to reach lower LOD and reduce the number of false positives rate during the diagnosis.

## RESULTS AND DISCUSSION

### DNA elements and type effect on protein expression

The ribosomal footprint and ribosome binding site (RBS) sequence have previously been shown to significantly impact protein synthesis, more than the promoter sequence, but in a more unpredictable way ^*51, 52*^. Even though the strength of the RBS relies heavily on the gene that is being translated due to mRNA structuring, a substantial amount of the currently provided parts characterizations are unapplicable for the substitution of genes ^*52*^. Therefore, computational modeling of DNA structure combined with experimental screening has been performed to find patterns in the DNA elements and expedite the design-build-test cycle for fast confirmation of the gene expression in the CFS. One of the popular computation tools is known as the RBS Calculator^*52, 53*^.

To see how well the computer-generated elements would predict protein synthesis, we only tested the highest *in silico* performing design of the 5’-UTR, RBS, and spacer region (ribosome footprint) and one output of the codon optimization for each of the fluorescent proteins. The RBS calculator has previously proven to be especially popular among *in vivo* protein expression studies ^*54*^, however here, we found the calculator does not fit to *in vitro* expression, even though the predicted translation initiation rate from the RBS calculator is significantly higher than the wild type (Figure 1A). We found that the predicted translation initiation rate (TIR) values were opposite from the actual expression for both sfGFP and mNG, testing the optimization of the RBS, codon sequence, and both combined. This is almost expected since inserting new elements into DNA expressed in vivo is nonconventional, but rather requires many variations until the desired function is reached ^*55*^. This discordance is elevated when the system is taken *in vitro* where the expression environment becomes even more non-native. Another lab also discovered that the Ribosome Binding Calculator was not suitable for their in vitro protein expression, showing the least out of the 5 they tested, but it had one of the highest RNA expression rates ^*51*^. They were more successful with screening a subset of randomly generated RBS structures lacking strong structural elements.

**Figure 1.**
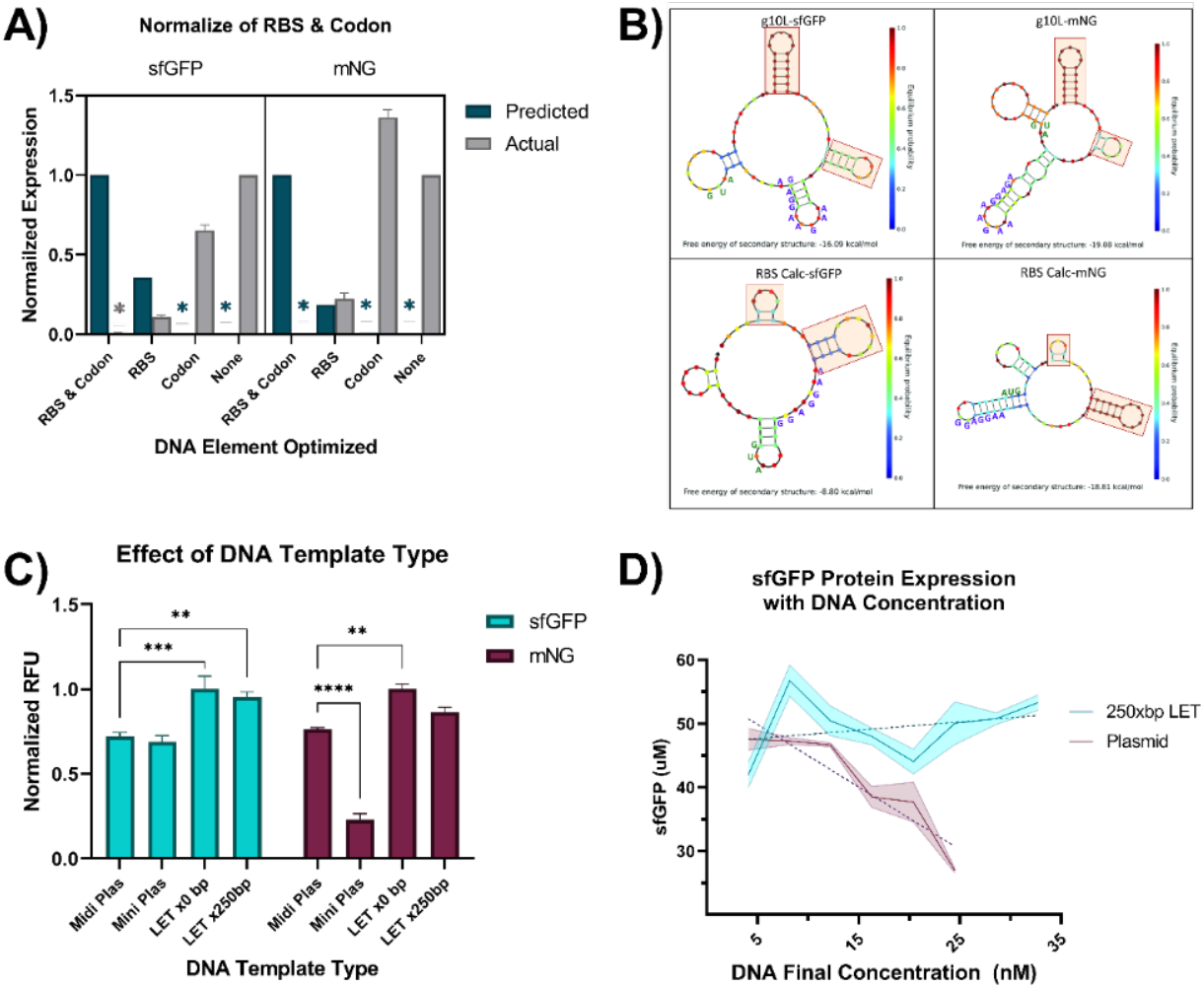
The effect of DNA elements and DNA template type on fluorescent protein output. **A)** RBS Calculator predicted TIR compared to actual fluorescent output (RFU) for the two fluorescent proteins. Colored “*” represent values too low to be seen on the graph. Predicted values were normalized to the highest value in each group. Actual values were normalized to the WT RFU expression in each group. All actual values for sfGFP and mNG were significantly different across DNA element optimization and compared against predicted values (P<0.0001, P<0.05 for mNG RBS predicted vs. actual, two-way ANOVA, Tukey). **B)** NUPACK RNA prediction drawings of the sequences upstream of the RBS first 20 nucleotides for the g10-L sequence used in our WT expression and the RBS calculator enhanced TIR sequence. Standby site highlighted in red and RBS site and start codon spelled out in blue and green, respectively. At the bottom of each structure, it says “Free energy of secondary structure” and the values are -16.09, -19.08, -8.08, and -18.08 kcal/mol, respectively. On the sides of the structure, it says “Equilibrium probability.” **C)** sfGFP and mNG were expressed in four different template types. RFU values normalized to highest expressed within protein group fluorescence output for miniprep plasmid showed the most variation between sfGFP and mNG, with mNG miniprep plasmid showing consistently significantly lower expression while sfGFP did not show significantly lower expression (****P<0.0001 two-way ANOVA, Tukey). LETx0bp showed slightly higher expression than the heavily purified Midiprep plasmid template (sfGFP ***P<0.001, mNG using two-way ANOVA, Tukey). LETx250bp showed higher expression than Midiprep plasmids only for sfGFP expression (**P<0.01 two-way ANOVA, Tukey). **D)** Comparing sfGFP expression with different DNA concentrations from a plasmid and a LET with 250bp buffer on each end. Linear regression correlation R^2^=0.0747 for LET and R^2^=0.8520 for plasmid with slopes significantly different from each other, P<0.01. Plasmid slope was significantly non-zero (P<0.01), but not the slope of LET (P>0.05). All experiments in figures **A, C, and D** were run to completion (20 hrs) in the same conditions, data represented as mean ± SD, n=3.

The protein expression with our original RBS is significantly higher than the predicted RBS because we use the T7 phage gene 10 leader RNA (*g10*-L), which is a ribosome footprint that dramatically increases the protein expression of foreign genes in *E. coli*. This gene 10 in the T7 phage codes for coat protein, which is made the most after infection, so it has to have an optimal 5’-UTR for the T7 RNA polymerase and *E. coli* ribosomes to overproduce foreign proteins ^*56*^. Therefore, our 5’-UTR might already be at its optimal sequence for transcription and translation.

Since mRNA is single-stranded, most times, the RBS is not available for the 30S ribosomal subunit to immediately bind to since the mRNA binds to itself in a secondary structure, so it must bind to a non-sequence specific region flanked by a stable hairpin and the hairpin containing the RBS and await when the hairpin with the RBS opens so it can slide into place using linear diffusion and bind. This waiting region is known as the standby site, the most geometrically accessible and requires the least amount of RNA unfolding ^*57*^. NUPACK is a computer software that can predict these mRNA secondary structures (detailed methods in the supplemental information). We analyzed the RNA structure of the 5’-UTR and the first 20 bp of the coding region for our wild-type genes and RBS optimized genes. The *g10*-L sequence has very prominent hairpins upstream of possible standby sites (highlighted in red), and then a low structure around the RBS site and start codon (spelled out in blue and green, respectively) (Figure 1B, top). The calculated RBS footprints with the first 100bp show a less prominent standby site and more structure around the RBS site (Figure 1B, bottom). The TIR is directly correlated to the amount of energy the ribosome must spend on its own, doing things such as weaving through tall hairpins, unfolding RNA, and ribosomal distortion—with the more expended, the less left for translation initiation ^*58*^. Possibly the structures of the *g10*-L are more favorable for conserving the ribosome’s initial energy than the calculated ones, especially in a more dilute environment (the CFS) compared to the whole cell, which could change the electrostatic interactions of the ribosome and RNA even more.

Not only does the RBS sequence make a difference in how well RNA is transcribed, but so does the codon sequence. Researchers have found that high GC content in the coding region creates more protein, and the cell’s codon usage can impact protein expression levels greatly by influencing the folding speed and efficiency of the protein during translation ^*51, 59, 60*^. This is mainly due to the charged tRNA pools, the cell’s usage of synonymous codons and rare codons in recombinant genes, which makes protein synthesis stall or perform incorrectly if the rare tRNA’s become depleted ^*61, 62*^. The benefits of codon optimization for recombinant protein synthesis in the CFS have been shown before, resulting in a 7.4-fold increase in protein for the cell extract void of rare tRNA expression ^*63*^. We found that codon optimizing increased mNG expression (x1.3 times) but decreased sfGFP expression (x0.8 times) (Figure 1A). Since mNG was not codon-optimized for *E. coli* codon usage and sfGFP was already established as a model reporter protein in *E. coli*, so sfGFP’s codon usage is possibly already well-coordinated to fit for *E. coli*.

The type of DNA template used in the CFS plays a crucial role in the speed of design-build-test (DBT) cycles. Previously, it has been demonstrated that linear DNA expression templates (LETs) amplified by PCR perform very well in the CFS when extra base pairs (bp) are added to protect the important DNA elements (5’- and 3’-UTRs) from any degradation at the ends, some showing 26-fold (mRFP1) and 12-fold (GFPmut3b) increase in expression ^*51*^, and 6-fold increase (deGFP) ^*64*^ from LETs with no buffer of base pairs. In our experiments, we did not see a significant increase in expression by adding 250 extra-base pairs to the 5’ and 3’ ends of the LET (upstream of the promoter and downstream of the terminator), but we observed a small increase in expression compared to the plasmid templates of the fluorescent proteins (Figure 1C). Even though the expression of LETs with the 250bp buffer was slightly lower than those without, we decided to use the 250bp buffer LETs from there on out to remove some possibility of gene degradation during storage. The exonuclease inhibitor GamS also can increase expression from linear DNA templates with 250bp buffer by 26% for deGFP, since it helps protect the ends from degradation ^*64*^. When a final concentration of 2uM of GamS was used in our experiments with linear DNA with 250bp buffer on the 5’ and 3’ ends, we saw a 14.0 ± 2.5 % increase in fluorescence for sfGFP and 24 ± 5.1 % increase for mNG (Figure S1). We also tested Miniprep-level purified plasmid DNA templates since they are much faster and less expensive to purify than plasmids purified at the midi and maxiprep levels. Interestingly, sfGFP expression was not affected by the lack of extra purification steps, but mNG was significantly affected, with a ∼70% decrease, even after repeating the experiment multiple times (Figure 1C).

The maximum concentration of plasmid DNA for the maximum amount of protein has been shown to plateau at 5-15 nM of DNA concentration, but the plateau for LET was not recorded since the experiment stopped at 15 nM ^*64, 65*^. Here we sought to find the plateau of a LET template and compare it with that of the plasmid’s (Figure 1D). Expressing sfGFP, we found that the plasmid plateau was lower than previous reports, possibly since we were expressing a different fluorescent protein (sfGFP) instead of deGFP, with their plateaus occurring around 5-10 nM and decreasing soon after. However, LET gene expression did not have a clear plateau, and expression was extremely variable across concentrations.

### mNeonGreen, a brighter fluorescent protein than those commonly used

In this paper, we debut the usage and production of mNG in the CFS for the first time, as an extremely bright reporter protein that can be used with CF sensors. mNG is not derived from the original green fluorescent protein variants from *Aequorea victoria*, but is derived from the cephalochordate *Branchiostoma lanceolatum*, and is the monomeric variant of the tetramer LanYFP ^*66*^. It is the brightest monomeric green or yellow fluorescent protein reported to date. It has a high quantum yield (∼0.80) and extinction coefficient (∼116,000 M^-1^cm^-1^). It has shown a high acid tolerance, with a pKa of 5.7, making it a good candidate for long-term expression in the CFS, which turns acidic as ATP is consumed ^*67*^.

In CF reactions, green fluorescent proteins provide the fastest response and lowest LOD compared to red fluorescent reporters and colorimetric LacZ output ^*68*^, so we compared the new mNG protein with the popular green proteins: sfGFP, deGFP, eGFP, and YPet in the CFS. The fluorescence characterizations of each protein can be found in Table S2. deGFP ^*65*^ was created to be more translatable in the CFS than eGFP, but here we show mNG is 7.75-fold brighter than deGFP and 26.81-fold brighter than eGFP, and even significantly brighter than the super-folder GFP, 2.08-fold, a commonly used protein for CF sensors due to its brightness (Figure 2A). mNG is not just brighter than the commonly used sfGFP because the system is making more of the protein. As shown in Figure 2B, the RFU/uM ratio of mNG is 2.1-fold higher than sfGFP, meaning the protein itself is brighter than sfGFP in our CFS. mNG was also compared to Ypet, a yellow fluorescent protein, and it showed to be brighter by 3.21-fold. The excitation and emission spectrums for all of the fluorescent proteins are shown in Figure 2C, with sfGFP, deGFP, and eGFP sharing the same curves.

**Figure 2.**
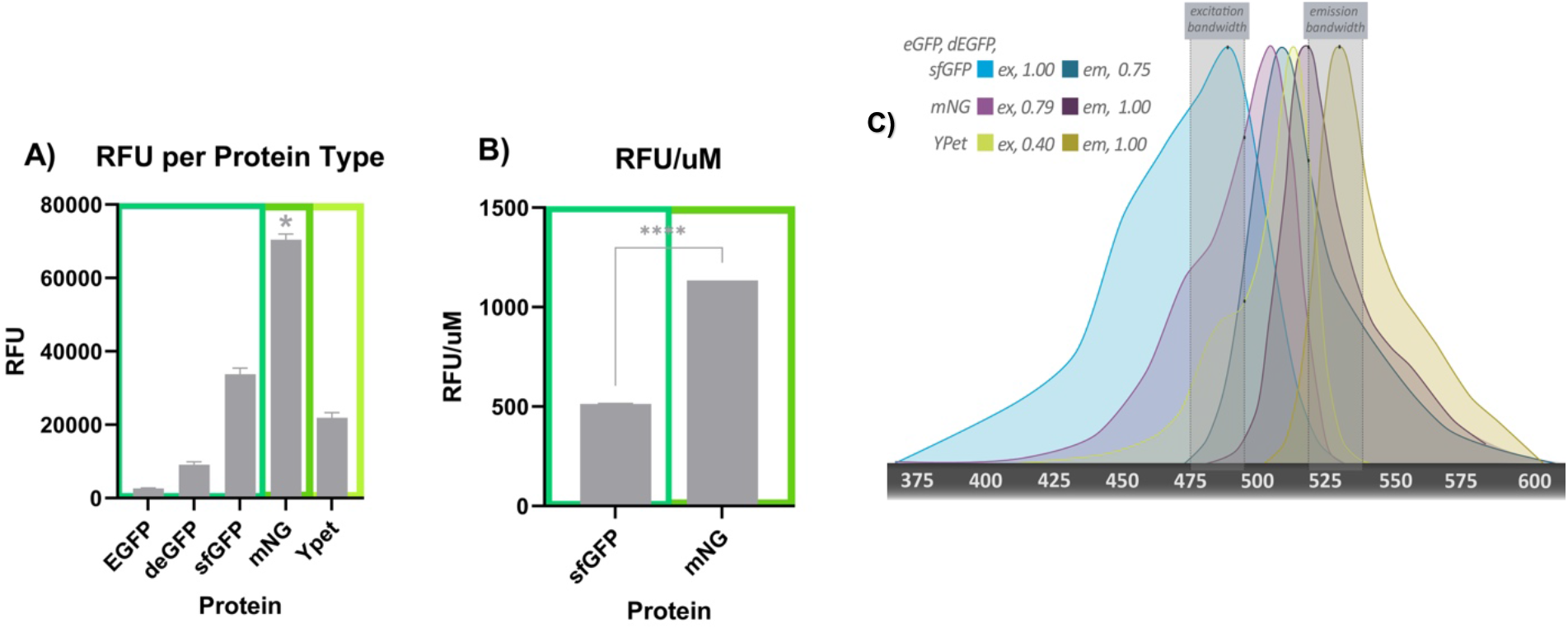
Comparing common CF fluorescent proteins. **A)** eGFP, deGFP, sfGFP, mNG, and YPet were synthesized in the CFS in LET DNA format and compared using the same filters. mNG was significantly brighter than all other fluorescent proteins tested (*P<0.0001 using one-way ANOVA), data represented as mean ± SD, n=3. **B)** RFU/μM comparison of sfGFP and mNG showing mNG is significantly brighter than sfGFP and not just making more protein, with an RFU/μM ratio 2.1-fold higher than sfGFP (****P<0.0001 using an Unpaired t-test), data represented as mean ± SD, n=3. **C)** Comparing the fluorescent excitation-emission spectrums of all 5 proteins. eGFP, deGFP, and sfGFP had extremely similar peaks so they were included as one (blue), mNG in purple, and YPet in yellow. The maximum relative intensities for excitation and emission of the proteins, given our filters, were mentioned in the top left corner for each protein and marked on the graph in black. The light white panels signify our filters’ bandwidths.

### Fluorescent output variability with environmental changes

Matrix effects ^*31-33*^ caused by non-native reagents and possible harmful molecules in the CFS sensor platform can unexpectedly interfere with the sensor’s function. These substances can range anywhere from buffers to samples stored in biological components. To measure the potential substance interference on the protein expression of the CFS sensor, we analyzed common additives, storage buffers, and environmental contaminants.

The extra molecular crowding by adding crowding agents and increased concentration of total proteins in the cell extract improve the overall cell-free protein synthesis level in general. To compare the fluorescent signal responsiveness in a liquid biopsy or storage additive condition, we measured the fluorescent output of sfGFP and mNG with 2% PEG8000 and increased cell extract concentration in the reaction mixture (26.7% to 40.0% v/v). Interestingly, sfGFP showed more evident signal flux at both crowding conditions, whereas mNG maintains consistent signal outputs (Figure 3A).

**Figure 3.**
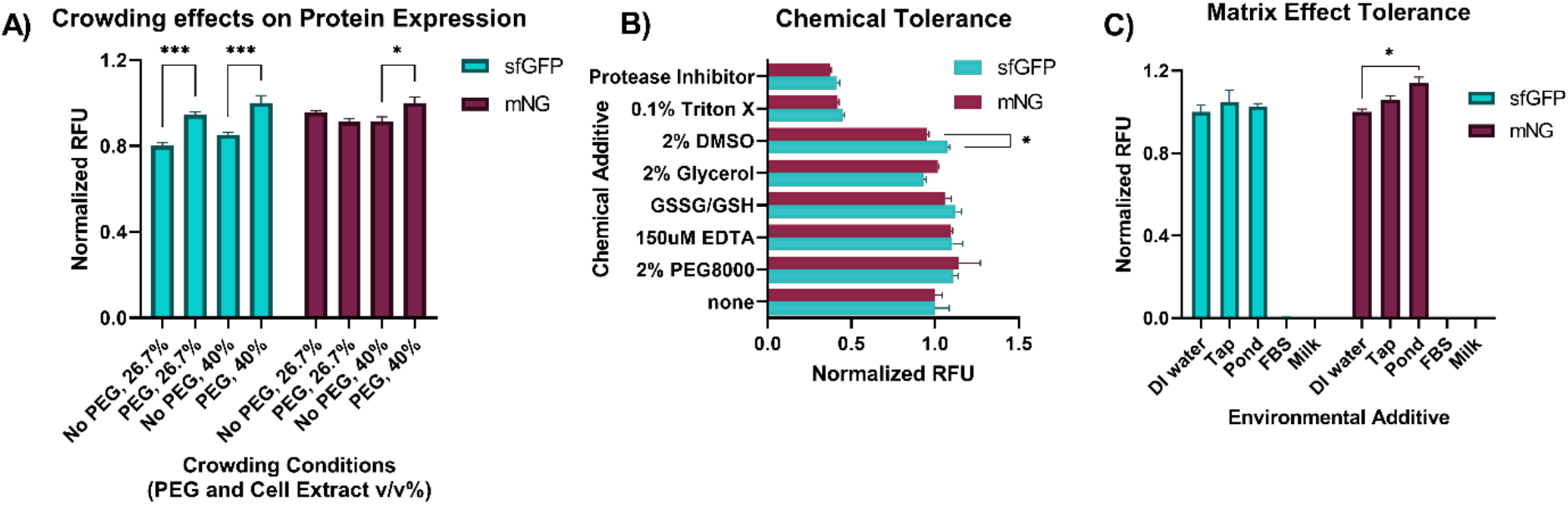
Fluorescent output and variability with environmental changes. **A)** Crowding effects on protein expression for sfGFP and mNG. Crowding was simulated by increasing the extract concentration and by adding 2% PEG8000. Expression was significantly increased for sfGFP in both extract concentrations and mNG for the higher extract concentration (*P<0.05, ***P<0.001 two-way ANOVA, Tukey). RFU values were normalized to the highest in each protein group separately. **B)** sfGFP and mNG tolerance to different chemical additives. sfGFP and mNG tolerated the chemical additives similarly, but sfGFP tolerated 2% DMSO more than mNG (*P<0.05, two-way ANOVA, Tukey). RFU values are normalized to the highest in each protein group separately. **C)** Comparing the ability of sfGFP and mNG to tolerate matrix effects with additions of 26.7% (v/v). Both could tolerate tap water and pond water well, with mNG even expressing significantly higher in pond water than DI water (*P<0.05, two-way ANOVA, Tukey). Protein synthesis did not occur with additions of FBS and whole milk. All data **A-C** represented as mean ± SD, n=3.

Next, we conducted the chemical tolerance test for sfGFP and mNG to compare the previous report ^*64*^ and matrix effect tolerance test. Both proteins tolerated the chemical additives and were not significantly different from each other (according to a two-way ANOVA), with the exceptions of 0.1% Triton X and Protease inhibitor, which reduced both proteins’ output signals by 55-60% (Figure 3B). Notably, Triton X and protease inhibitors are frequently used for the eukaryotic cell lysis process and can potentially interrupt the output signals when sensing specimens from the eukaryotic cells. We introduced common environmental additives, including tap water, pond water, whole milk, and fetal bovine serum, to the system that might contain an analyte of interest in the future. Previous research had shown that the CFS could perform in the presence of RNase A when RNase inhibitor presented. Since RNase inhibitor is costly, we chose to use polyvinylsulfonic acid (PVSA) which has been shown to mimic the functions of commercially available RNase inhibitors in the CFS with inhibiting RNases ^*69*^. PVSA was added to all reactions containing an environmental additive. Unfortunately, we did not obtain the same results when we used cell-culture media designated fetal bovine serum (FBS) which is a sample abundant with RNases and closer to a real-world application sample. Another study has added human serum to the CFS enabled the synthesis of proteins with Murine RNase inhibitor and only added 14% final volume fraction of serum to the reaction, while we added 26% with PVSA ^*70*^. Interestingly, pond water and tap water performed very well in the CFS. However, FBS and milk additives suppressed the output signals completely (Figure 3C).

### System optimization for mNG

In the cell-free protein synthesis reaction, many components are added, some having a higher impact on the protein synthesis outcome than others and some needing personalization for the specific protein being made ^*63, 71*^. Since it had not been synthesized in this system before, we screened mNG expression with different CFPS conditions, including pH, Mg^2+^ concentration, cell extract concentration, reaction temperature, and combinations of these mentioned.

To change the pH of the system, we used HEPES buffer with different pH’s. HEPES is used to stabilize the system’s pH, so changing this can have a big impact on the final pH of the system (Table S3) ^*71*^. We first used 250xbp LETs of mNG and sfGFP in these different pH environments but the trend was not consistent (Figure 4A). We then switched to using plasmids to express the proteins and noticed a more consistent trend, with the HEPES buffer (pH 7.8) showing the highest expression of mNG (Figure 4A). We concluded expression from LETs could vary much more across pHs than plasmids, therefore we used plasmids for the remainder of the experiments. Data for sfGFP was not shown but had the same trend. mNG and sfGFP expression levels were characterized at different final Mg^2+^ concentrations (4-18 mM), showing a similar trend for both but with sfGFP producing its most at 6 mM and mNG at 8 mM, respectively (Figure 4B). Then, mNG expression was tested in the combination of Mg^2+^ concentrations (8-12 mM) and pHs (6.9- 7.8). The pattern of low to high expression with the rise in pH was similar across Mg^2+^ concentrations, as well as a high expression with decreasing Mg^2+^, except for 10 mM Mg^2+^ and pH 6.9 (Figure 4C).

**Figure 4.**
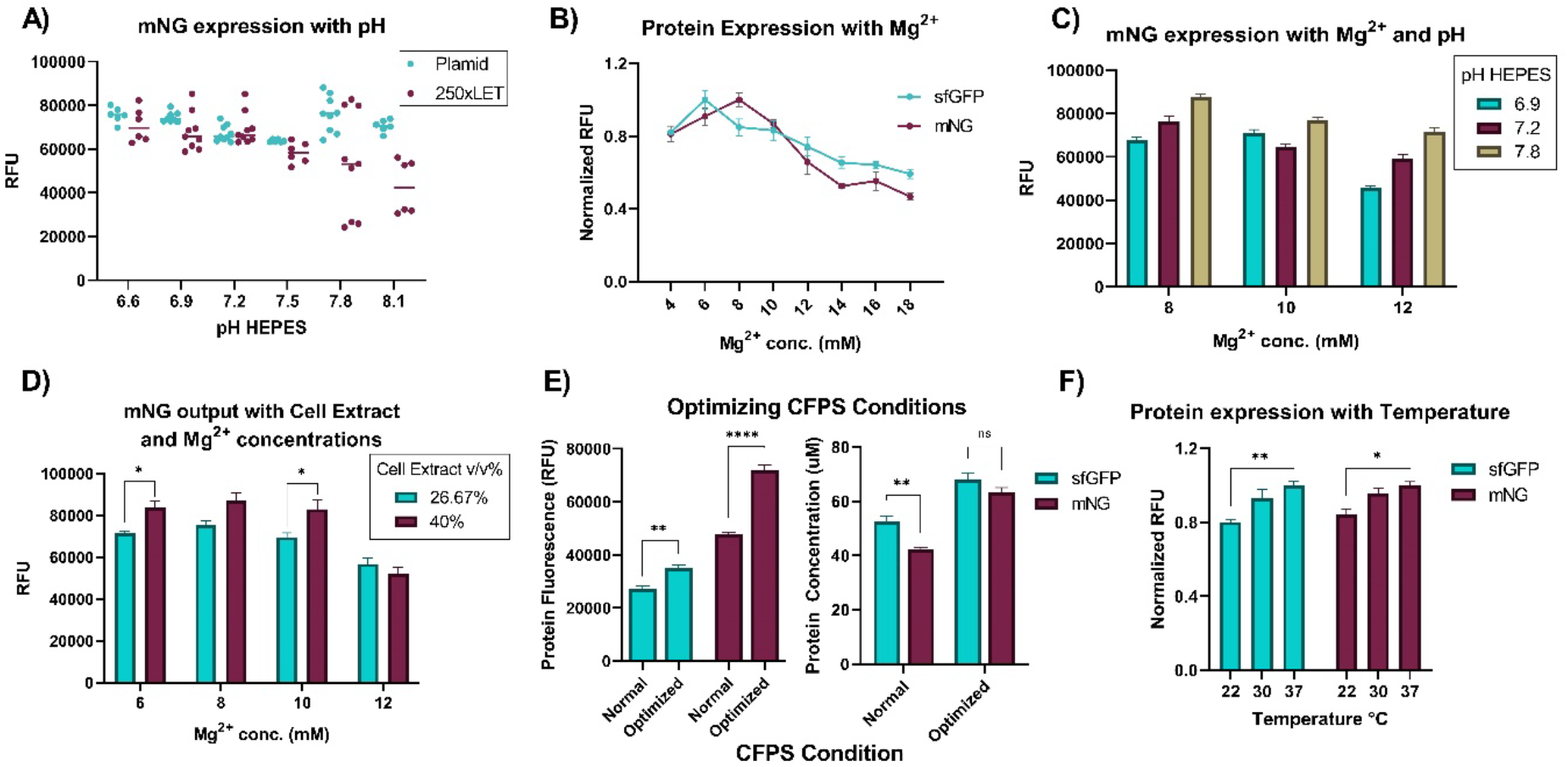
Characterizing protein expression and optimizing conditions. **A)** mNG expression with different pH’s of the buffer varies depending on the DNA template type. LET show more variability than plasmid DNA gene expression. HEPES buffer of pH 7.8 shows the highest expression. Data represented as mean ± SD, n=6-9. **B)** mNG and sfGFP expression with varying Mg^2+^ concentrations. Both show similar patterns, but with sfGFP producing its most at 6 mM and mNG at 8 mM. Data represented as mean ± SD, n=3. **C)** mNG expression with different Mg^2+^ concentrations and HEPES pH in the CFPS reaction. The pattern of low to high expression with the rise in pH was similar across Mg^2+^ concentrations, as well as a high expression with decreasing Mg^2+^, except for 10 mM pH 6.9. Data represented as mean ± SD, n=3. **D)** mNG expression with varying Mg^2+^ concentrations and cell extract v/v%. Adding more cell extract increases expression significantly in Mg^2+^ concentrations 6 mM and 10 mM (*P<0.05 two-way ANOVA, Tukey). Data represented as mean ± SD, n=3. **E)** mNG and sfGFP expression in optimized conditions compared to normal. The RFU value is shown on the right and protein concentration (μM) on the left. mNG has a higher RFU/μM ratio (Figure 2B), therefore the μM was slightly lower than sfGFP in optimized conditions, while significantly lower in the normal conditions, (**P<0.01 two-way ANOVA, Tukey). Data represented as mean ± SD, n=4. Fluorescence captured at a lower gain here. **F)** mNG and sfGFP expression with different temperatures. sfGFP expression is more significantly lowered at room temperature from 37 °C than mNG (*P<0.05, **P<0.01 two-way ANOVA, Tukey). RFU values were normalized to the highest in each protein group separately. Data represented as mean ± SD, n=3.

Since the Buffer B (S30 buffer) of the cell extract contributes to the final concentration of Mg^2+^ in the cell-free reaction, we tested mNG expression in different concentrations of cell extract across varying Mg^2+^ concentrations. The extra Mg^2+^ from the Buffer B did not influence the pattern of mNG synthesis across Mg^2+^ concentrations but adding more extract did improve expression significantly in Mg^2+^ concentrations 6 mM and 10 mM, and slightly in concentration 8 mM (Figure 4D). After the optimal conditions of pH, Mg^2+^, and cell extract concentration were verified, we then combined them into one experiment and compared them with the previously optimized CFPS (normal) conditions of our lab. These optimal conditions included a HEPES buffer (pH 7.8), 40% v/v cell extract, 2% PEG 8000, and 6 mM (mNG) and 8 mM (sfGFP) Mg^2+^. This is compared to the previously used conditions: HEPES buffer (pH 7.2), 26.7% v/v cell extract, no PEG 8000, and 12 mM Mg^2+^. These optimal CFPS conditions were able to increase fluorescence outputs by 50.3% and 28.4% for mNG and sfGFP, respectively (Figure 4E, left). The amount of protein synthesized in the normal conditions was 42.32 ± 1.22 uM or 1.13 ± 0.03 mg/mL for mNG and 52.41 ± 4.3 uM or 1.41 ± 0.12 mg/mL for sfGFP. The amount of protein synthesized in the optimal conditions was 63.33 ± 3.39 uM or 1.69 ± 0.09 mg/mL for mNG and 68.05± 5.07 uM or 1.83 ± 0.14 mg/mL for sfGFP (Figure 4E, right). Notably, mNG showed brighter signal output with a higher RFU/uM or mg/mL ratio. Therefore its protein concentration will be lower than sfGFP even though the RFUs are high. Lastly, protein synthesis was tested at different temperatures (22 °C (room temperature), 30 °C, and 37 °C) to compare the fluorescent signal outputs at various sensing temperatures. Both proteins had a slight drop in protein expression at room temperature, but mNG was able to tolerate it slightly more than sfGFP, making it a good candidate for a point of care reporter protein (Figure 4F). Throughout the optimization process, from sfGFP expression at normal conditions to mNG at optimal conditions, fluorescence readout was able to be increased 2.64-fold, which is a significant increase to aid in detecting low levels of analyte in the CFS while still maintaining a distinguishable readout.

### Maturation and fluorescent output speed

The maturation process of the GFP family involves the folding of the β-barrel, torsional rearrangements, cyclization, and oxidation and dehydration of the chromophore ^*72*^. The “super- folder” GFP compared in this study was engineered to fold more robustly and faster with more stability than the regular reporter GFP. These enhancements contribute to a generation of higher fluorescence signal outputs ^*50*^. We compared the protein maturation rate with the actual fluorescent signal outputs to apply mNG as an alternative fluorescent protein for the CF sensor. mNG has a comparable maturation to sfGFP in the CFS when measuring to a certain fold change, but mNG then keeps maturing to become even brighter (Figure 5A). sfGFP shows a 4-fold higher maturation rate k=26.62 × 10^−3^ s^-1^ and a 4-fold shorter half-time of maturation (t_1/2_) is 26.04 min when compared to mNG rate k=6.66 × 10^−3^ s^-1^ and half-time t_1/2_=104.7, but mNG shows a 2.32-fold greater fold change (4.322 ± 0.846) after the translation has ended than sfGFP (1.867 ± 0.370). This maturation experiment was conducted by letting the reaction run for 75 minutes and then immediately putting the tubes on an ice slushy for 5 minutes. Then, the tubes were stored at 4 °C. The rate of the fluorescent output showed that the fluorescent proteins fluoresced about 10 mins apart from each other, with sfGFP in the lead (Figure 5B). Although, mNG fluorescence fits more of an exponential pattern with a higher R^2^ value (0.9394) than sfGFP (R^2^= 0.8692). They are both able to be visualized by the naked eye at around the 45-minute mark when DNA template concentrations are close to saturation points in a 15 µL reaction. Overall, both proteins express fluorescence promptly in the CFS.

**Figure 5.**
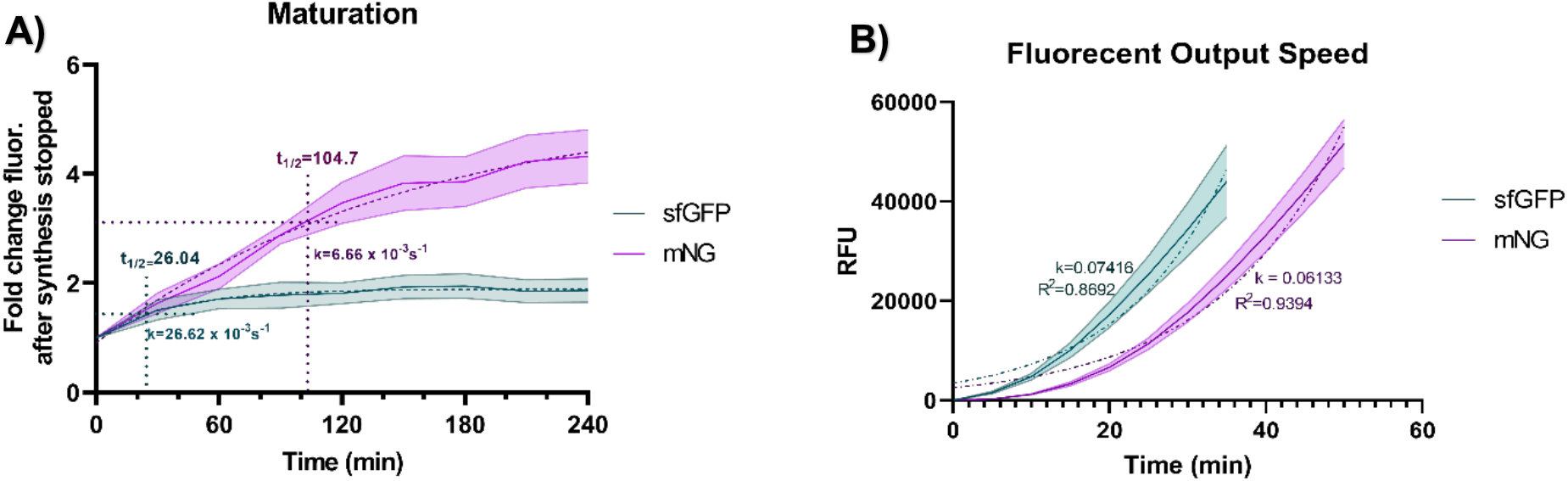
Fluorescence kinetics of sfGFP and mNG. **A)** Maturation of sfGFP and mNG. Fluorescence fold change after synthesis had ceased. sfGFP maturation rate k=26.62 × 10^−3^ s^-1^ and half-time of maturation (t_1/2_) 26.04 min. mNG maturation rate k=6.66 × 10^−3^ s^-1^ and half-time of maturation t_1/2_=104.7. **B)** Fluorescent output speed of sfGFP and mNG. sfGFP exponential growth rate k=0.07416 and mNG k=0.06133. RFU was measured on a different machine than in the other experiments. **A and B** are represented as mean ± SD, n=3.

## CONCLUSION

In this research, we improved the fluorescence output of mNG in the CFS by 2.64-fold when compared to sfGFP and characterized protein synthesis in different conditions to offer mNG as an alternative fluorescent protein to achieve a low LOD. This was accomplished by first introducing a brighter fluorescent protein, mNeonGreen. Next, we identified more favorable DNA settings in the CFS with codon optimization, clarifying the optimal RBS, 5’-UTR and spacer sequence, and finding the optimal DNA template and type. We also characterized the fluorescent signal outputs of mNG and sfGFP while enduring different matrix effects from biological samples and possible CFS additives. Comparing mNG to the “super-folder” sfGFP, we found mNG matured slower than sfGFP, but reached the fluorescent maturation plateau point at the same time and surpassed it with a 2.32-fold greater change in maturation. Lastly, we optimized reaction conditions through a systemic optimization process of Mg^2+^ concentration, pH, percentage of cell extract, and molecular crowding conditions. In conclusion, these combinations can be used to increase the dynamic range of a CF sensor by optimizing the fluorescent signal output itself instead of amplifying the target analyte so sensors can reach a lower LOD while keeping the number of false-positives low.

## METHODS

### Materials

*Escherichia coli strains* Subcloning Efficiency™ *DH5α* [Genotype F^−^ Φ80*lac*ZΔM15Δ(*lac*ZYA-*arg*F) U169 *rec*A1 *end*A1 *hsd*R17(r_K_^−^, m_K_^+^) *pho*A *sup*E44 *thi*- 1 *gyr*A96 *rel*A1 λ-] and BL21 star (DE3) [Genotype F^−^ *ompT hsdS*_B_ (r_B_^−^m_B_^−^) *gal dcm rne131* (DE3)] were used for plasmid cloning and a source of the cell extract, respectively (Invitrogen, Waltham, MA). The *E. coli* cells were grown in either Luria–Bertani (LB) media (10 g/L of tryptone, 5 g/L of yeast extract, 10 g/L of sodium chloride in Milli-Q water) or 2xYTPG media (16 g/L of tryptone, 10 g/L of yeast extract, 5 g/L of sodium chloride, 7 g/L of potassium phosphate dibasic, 3 g/L of potassium phosphate monobasic, pH 7.2, and 0.1 M of glucose in Milli-Q water). CFS components including *E. coli* total tRNA mixture (from strain MRE600) ATP, GTP, CTP, UTP, Phosphoenolpyruvate, 20 amino acids, and other materials were purchased from Sigma-Aldrich (St. Louis, MO), Alfa Aesar (Haverhill, MA), and Fisher Scientific (Hampton, NH),

### DNA preparation

The optimal RBS for every gene was found using the De Novo DNA web tool known as the RBS Calculator ^*52*^. Primer extension was used to add different RBS footprint sequences to sfGFP and mNG. The codon-optimized versions of sfGFP and mNG sequences were prepared using the codon optimization tool for *E. coli* codon usage bias (strain K12) (Integrated DNA Technologies, Coralville, IA) and inserted into pJL1 plasmid using Gibson Assembly. Plasmids were transformed into DH5α electrocompetent cells for cloning and purification (E.Z.N.A.® Plasmid DNA Midi Kit, Omega Bio-tek, Norcross, GA) for sequencing and cell-free reaction, respectively.

Linear templates were prepared by PCR and subsequent purification (E.Z.N.A.® Cycle Pure Kit, Omega Bio-tek, Norcross, GA). The oligomer sequences are listed in Table S1. The amplified region included buffer sequences (250 bp each at 5’- and 3’-ends) upstream of the promoter and downstream of the terminator sequences unless otherwise noted (Table S1). The PCR products were verified on a 1% agarose to confirm the size and low off-target amplification. The DNA concentrations were measured with the BioTek Synergy HTX multi-mode reader using Take3 Micro-Volume Plate.

### Preparation of cell extract

Cell extract was prepared as described previously ^*73, 74*^. Briefly, overnight cultured *E. coli* BL21 Star (DE3) in LB media was inoculated to sterilized 1 L 2xYTPG media in a 2.5-L baffled Tunair shake flask and the cells were cultured at 37°C with vigorous shaking at 250 rpm. The optical density of the cells was monitored by the UV-Vis spectrophotometer (Genesys 6, Thermo Fisher Scientific, Waltham, WA) (Figure S2) and induced to overexpress T7 RNA polymerase at OD_600_ 0.6 with 1 mM of isopropyl β-D-1-thiogalactopyranoside (IPTG) (Figure S2 and S3). Cells were harvested at the mid-exponential phase (OD_600_ 2.0) by centrifugation (5,000 RCF at 4 °C for 15 min). Cell pellets were then washed with Buffer A (DTT added, pH 7.8) three times and flash-frozen in liquid nitrogen and stored at a -80°C freezer until use. Cell pellets were used within the same week to ensure maximum activity. Cell pellets were thawed on ice and then resuspended with Buffer B (no DTT, pH 8.2) with 1 mL of Buffer B for 1 g of wet cell mass and transferred into microtubes in 1 mL aliquots for lysis. The sonicator (Q125, Qsonica, Newtown, CT) with a 1/8” diameter probe was set to 20-kHz frequency, 50% amplitude, and 10 s on and 10 s off. To minimize protein degradation by heat, the tubes of cells were kept in ice-water during sonication. The number of joules was determined by the equation found previously ^*74*^, which equates to 537 J for 1 mL of resuspended cells. 3 µL of 1 M DTT was added per 1 mL lysate. The tubes were then centrifuged at 12,000 RCF at 4 °C for 10 minutes and the supernatant was taken. Run-off and dialysis were not performed for the experiments in this study as we found them to be inhibitory, especially for LET DNA (Figure S4). The cell extract used in CFPS reactions, was aliquoted and flash-frozen in liquid nitrogen and stored at a -80 °C freezer until use. The extract aliquots were only thawed for the reaction they were used for and not reused again to ensure the activity.

### Cell-free protein synthesis

Cell-free protein synthesis reactions were carried out in 1.5 mL microtubes in an incubator. The reaction volume was 15 μL with the following components: 1.2 mM ATP; 0.85 mM each of GTP, UTP and CTP; 34.0 μg/mL L-5-formyl-5, 6, 7, 8-tetrahydrofolic acid (folinic acid); 170.0 μg/mL of *E. coli* tRNA mixture; 130 mM potassium glutamate; 10 mM ammonium glutamate; 12 mM magnesium glutamate; 2 mM each of 20 amino acids; 57 mM of HEPES buffer (pH 7.2, except for the pH experiments in Figure 4); 0.4 mM of nicotinamide adenine dinucleotide (NAD); 0.27 mM of coenzyme A; 4 mM of sodium oxalate; 1 mM of putrescine; 1.5 mM of spermidine; 33 mM phosphoenolpyruvate (PEP); and 27% v/v of cell extract. DNA was added at 13.3 μg/mL for plasmid DNA. Linear DNA final concentrations varied depending on length but were around 4.95 μg/mL for LETs without buffer sequences and 7.58 μg/mL for LETs with 250 bp buffer sequences. For the matrix effect experiments, samples were added at 26.67% of the final volume. Reactions were run for 20 hours to ensure completion at 30 °C unless otherwise mentioned.

### Quantitative analysis of fluorescent proteins

The fluorescence intensity of the synthesized fluorescent proteins was measured by the multi-well plate fluorometer (Synergy HTX, BioTek, Winooski, VT). 5 µL of the cell-free synthesized fluorescent protein and 45 µL of Milli-Q water were mixed in a 96-well half-area black plate (Corning Incorporated, Corning, NY). The plate was mixed in the plated reader orbitally at medium speed for 15 s and read at the height of 1.5 mm with a gain of 48. The excitation and emission spectra are 485 and 528 nm, respectively. The cell-free synthesized protein was visualized by Coomassie blue staining after protein gel electrophoresis using pre-casted 4-12% Bis-Tris gradient gel (Invitrogen, Waltham, MA) (Figure S5).

### Statistical analysis

Statistical analyses were conducted using Graphpad Prism 8.4.3 (GraphPad Software) with a 5% significance level. For the parametric analysis of data from quantification of the synthesized protein, two-way ANOVA followed by the Dunnett’s test was used.

## AUTHOR CONTRIBUTION

C.E.C. and Y.-C.K. conceived the project. C.E.C., J.K., and Y.-C.K. designed and conceptualized experiments. C.E.C., P.L.C., and C.J.H. prepared and performed experiments and acquired data. C.E.C. and J.K. analyzed and interpreted data. C.E.C. wrote the original manuscript. C.E.C., J.K., and Y.-C.K. revised and edited the manuscript. All authors contributed to the article and approved the submitted version.

## ACKNOWLEDGEMENT

This work was supported by Louisiana Board of Regent (RCS, Grant No. LEQSF(2020- 23)RD-A-01) and USDA National Institute of Food and Agriculture (HATCH, Accession No. 1021535, Project No. LAB94414).

