## Supplemental Information for "Characterizing a New Fluorescent Protein for Low Limit of Detection Sensing in the Cell-Free System"

### Table S1. DNA sequences (5’ to 3’) used in this study

| | |
|---|---|
| T7 promoter | taatacgactcactatagg |
| T7 terminator | ctagcataaccccttggggcctctaaacgggtcttgaggggttttttg |
| 5’-UTR g10-leader, RBS, & spacer | gagaccacaacggtttccctctagaaataattttgtttaactttaagaaggagatatacat |
| sfGFP RBS calculator 5’-UTR, RBS, & spacer | gagaccaaacggaacagtcggtacggagattaaggaggtttaa |
| mNG RBS calculator 5’-UTR, RBS, & spacer | gagaccataactcgggatcatattcccgacatttttctataaggaggttttttt |
| sfGFP WT ORF & *strep tag* | atgatgagcaaaggtgaagaactgtttaccggcgttgtgccgattctggtggaactggatggcgatgtgaacggtcacaaattcagcgt<br>gcgtggtgaaggtgaaggcgatgccacgattggcaaactgacgctgaaatttatctgcaccaccggcaaactgccggtgccgtggcc<br>gacgctggtgaccaccctgacctatggcgttcagtgttttagtcgctatccggatcacatgaaacgtcacgatttctttaaatctgcaatgc<br>cggaaggctatgtgcaggaacgtacgattagctttaaagatgatggcaaatataaaacgcgcgccgttgtgaaatttgaaggcgatacc<br>ctggtgaaccgcattgaactgaaaggcacggattttaaagaagatggcaatatcctgggccataaactggaatacaactttaatagccat<br>aatgtttatattacggcggataaacagaaaaatggcatcaaagcgaattttaccgttcgccataacgttgaagatggcagtgtgcagctg<br>gcagatcattatcagcagaataccccgattggtgatggtccggtgctgctgccggataatcattatctgagcacgcagaccgttctgtct<br>aaagatccgaacgaaaaacgggaccacatggttctgcacgaatatgtgaatgcggcaggtattacg*tggtcccatccgcagtttgaa*<br>*aag*taataa |
| sfGFP codon optimized ORF & *strep tag* | atgagtaagggtgaggagcttttcacaggggtagtcccgattcttgtggagttagatggcgacgtaaacggccataaattctctgtgcg<br>gggggaaggtgaaggcgacgcgacaattggcaaacttactctcaagtttatctgcacaaccgggaagctcccagtaccgtggccaac<br>gcttgtaacgacactgacatacggggttcagtgtttctcccgctacccggaccatatgaaacggcatgacttcttcaagagcgcaatgc<br>cggaaggctacgttcaggaacggaccatttcctttaaagatgatgggaaatataagactcgcgcggtagtaaaatttgagggcgacac<br>attggtaaaccggattgaactcaaaggcacagactttaaggaagatggcaacattctcgggcacaagttggaatataattttaattcgca<br>caacgtttatattaccgcagataaacagaaaaatgggatcaaggcgaacttcacagtacgccacaacgttgaggatggttctgtacaac<br>ttgctgaccattatcaacagaacacgcctattggtgatgggcctgtccttcttccagataatcactatttatcaacgcagaccgtcttgtcca<br>aagacccaaatgaaaagcgtgaccatatggtgttgcacgaatacgttaacgcagcgggcatcacc*tggtcccatccgcagtttgaaa*<br>*ag*taataa |
| mNG WT ORF | atggctagtttgcctgctacccacgaattacacatttttggatcaatcaatggagtcgattttgatatggttggtcaaggaaccggtaatcc<br>aaatgatggatacgaagaattgaatcttaagagtactaagggtgatttacagttttctccatggattttggtccctcatatcggatacggtttt<br>caccaatacttgccatatcctgatggaatgtctccttttcaggctgcaatggttgatggatcaggttatcaagtccatcgtacaatgcagttt<br>gaagatggtgctagtcttaccgttaattatcgttacacatatgaaggatctcacattaaaggtgaagcccaagtaaagggaacaggttttc<br>cagcagatggacctgttatgactaattcacttaccgccgctgattggtgtcgtagtaaaaagacttatccaaatgataagacaatcatctct |

| | |
|---|---|
| | acttttaagtggtcatacacaactggaaatggtaaacgttatcgttcaactgctcgtaccacatacacctttgcaaaaccaatggcagcca<br>attaccttaagaatcaacctatgtacgtctttcgtaagacagaacttaagcattctaagactgaacttaattttaaggaatggcagaaggctt<br>ttactgatgtaatgggtatggatgaattgtataaataa |
| mNG codon optimized ORF & *strep tag* | atggcaagtctacccgctacacacgaattacacatcttcggtagtattaacggggtggattttgatatggttggtcagggtactggaaacc<br>cgaatgacggctatgaggaactgaacctgaagtccaccaaaggtgatctgcaattctctccgtggattctggttccgcatatcggttacg<br>gcttccatcaatatttaccgtatccagatggcatgagcccatttcaggcggccatggtcgacggctctggttaccaagtgcatagaacca<br>tgcagttcgaggacggcgcgagcctgacggtgaactaccgctacacctacgagggttcccacatcaaaggcgaagcgcaggtgaa<br>aggtactggcttcccggcagacggtccggttatgaccaatagcttgaccgcggctgactggtgccgttcgaagaagacgtacccgaat<br>gacaaaaccattatctccaccttcaagtggagctataccaccggcaacggtaaacgttaccgcagcactgcgcgtaccacatacacctt<br>cgccaaaccgatggcagctaattatttgaagaaccagccgatgtatgtctttcgtaaaacggagcttaagcacagcaagaccgagctc<br>aactttaaagaatggcaaaaggcgtttaccgatgttatgggtatggatgaactgtataaa*tggtcccatccgcagtttgaaaag*taataa |
| T7 terminator 3’-UTR | gtcgaccggctgctaacaaagcccgaaaggaagctgagttggctgctgccaccgctgagcaataa |
| 250xbp LET 5’ buffer | cgaactgagatacctacagcgtgagcattgagaaagcgccacgcttcccgaagggagaaaggcggacaggtatccggtaagcggc<br>agggtcggaacaggagagcgcacgagggagcttccagggggaaacgcctggtatctttatagtcctgtcgggtttcgccacctctga<br>cttgagcgtcgatttttgtgatgctcgtcaggggggcggagcctatggaaacgaattcagatctcgatcccgcgaaat |
| 250xbp LET 3’ buffer | ctgaaagccaattctgattagaaaaactcatcgagcatcaaatgaaactgcaatttattcatatcaggattatcaataccatatttttgaaaa<br>agccgtttctgtaatgaaggagaaaactcaccgaggcagttccataggatggcaagatcctggtatcggtctgcgattccgactcgtcc<br>aacatcaatacaacctattaatttcccctcgtcaaaaataaggttatcaagtgagaaatcaccatgagtg |
| pJL1 vector backbone with Kanamycin resistance gene and origin of replication, ColE1 | ctgaaagccaattctgattagaaaaactcatcgagcatcaaatgaaactgcaatttattcatatcaggattatcaataccatatttttgaaaa<br>agccgtttctgtaatgaaggagaaaactcaccgaggcagttccataggatggcaagatcctggtatcggtctgcgattccgactcgtcc<br>aacatcaatacaacctattaatttcccctcgtcaaaaataaggttatcaagtgagaaatcaccatgagtgacgactgaatccggtgagaat<br>ggcaaaagcttatgcatttctttccagacttgttcaacaggccagccattacgctcgtcatcaaaatcactcgcatcaaccaaaccgttatt<br>cattcgtgattgcgcctgagcgagacgaaatacgcgatcgctgttaaaaggacaattacaaacaggaatcgaatgcaaccggcgcag<br>gaacactgccagcgcatcaacaatattttcacctgaatcaggatattcttctaatacctggaatgctgttttcccggggatcgcagtggtg<br>agtaaccatgcatcatcaggagtacggataaaatgcttgatggtcggaagaggcataaattccgtcagccagtttagtctgaccatctca<br>tctgtaacatcattggcaacgctacctttgccatgtttcagaaacaactctggcgcatcgggcttcccatacaatcgatagattgtcgcac<br>ctgattgcccgacattatcgcgagcccatttatacccatataaatcagcatccatgttggaatttaatcgcggcttcgagcaagacgtttcc<br>cgttgaatatggctcataacaccccttgtattactgtttatgtaagcagacagttttattgttcatgatgatatatttttatcttgtgcaatgtaac<br>atcagagattttgagacacaacgtggatcctgcagttgagatcctttttttctgcgcgtaatctgctgcttgcaaacaaaaaaaccaccgct<br>accagcggtggtttgtttgccggatcaagagctaccaactctttttccgaaggtaactggcttcagcagagcgcagataccaaatactgt<br>ccttctagtgtagccgtagttaggccaccacttcaagaactctgtagcaccgcctacatacctcgctctgctaatcctgttaccagtggct<br>gctgccagtggcgataagtcgtgtcttaccgggttggactcaagacgatagttaccggataaggcgcagcggtcgggctgaacggg<br>gggttcgtgcacacagcccagcttggagcgaacgacctacaccgaactgagatacctacagcgtgagcattgagaaagcgccacgc<br>ttcccgaagggagaaaggcggacaggtatccggtaagcggcagggtcggaacaggagagcgcacgagggagcttccaggggga<br>aacgcctggtatctttatagtcctgtcgggtttcgccacctctgacttgagcgtcgatttttgtgatgctcgtcaggggggcggagcctatg<br>gaaacgaattcagatctcgatcccgcgaaat |

| | |
|---|---|
| T715up FW (LET x0bp buffer) primer | tcgatcccgcgaaattaatacgactcactatag |
| T7 Terminator RV (LET x0bp buffer) primer | caaaaaacccctcaagacccgttta |
| pJL1x250bp LET FW primer | ccacctctgacttgagcgtcgat |
| pJL1x250bp LET RV primer | gcagcagccaactcagcttcctt |

#### Table S2. Characteristics of fluorescent proteins

| | Excitation λ | Emission λ | Extinction Coefficient | Quantum Yield | Brightness | pKa | Maturation (min) | Lifetime (ns) | Citation |
|---|---|---|---|---|---|---|---|---|---|
| eGFP | 488 | 507 | 55,900 | 0.6 | 33.54 | 6.0 | 25.0 | 2.6 | [1] |
| deGFP | 488 | 507 | * | * | * | 6.0 | * | * | [2] |
| sfGFP | 485 | 510 | 83,300 | 0.65 | 54.15 | 5.9 | 13.6 | * | [3] |
| mNG | 506 | 517 | 116,000 | 0.8 | 92.8 | 5.7 | 10.0 | 3.1 | [4] |
| Ypet | 517 | 530 | 104,000 | 0.77 | 80.08 | 5.63 | * | * | [5] |

*Some information could not be found

#### Table S3. HEPES pH vs. reaction pH before and after

| **HEPES** | **6.6** | **6.9** | **7.2** | **7.5** | **7.8** | **8.1** |
|---|---|---|---|---|---|---|
| Plasmid RXN before | 6.98 | 7.2 | 7.34 | 7.47 | 7.69 | 7.79 |
| Plasmid RXN after | 6.59 | 6.68 | 6.62 | 6.68 | 7.01 | 6.86 |
| **Difference:** | -0.39 | -0.52 | -0.72 | -0.79 | -0.68 | -0.93 |
| LET RXN before | 7.04 | 7.24 | 7.35 | 7.47 | 7.73 | 7.778 |
| LET RXN after | 6.92 | 6.99 | 7.01 | 7.12 | 7.36 | 7.25 |
| **Difference:** | -0.12 | -0.25 | -0.34 | -0.35 | -0.37 | -0.528 |

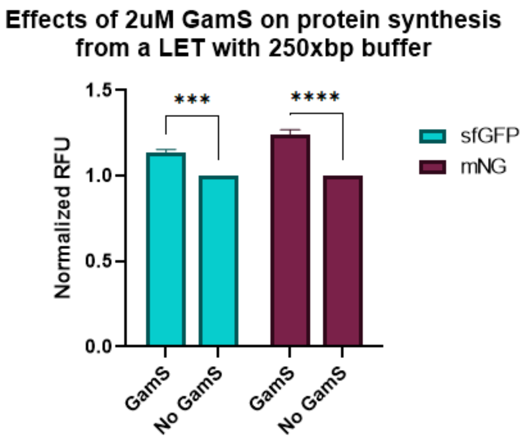


**Figure S1. Effects of 2 µM of GamS on protein synthesis for a LET with 250xbp buffer.** (***P<0.001, ****P<0.0001, two-way ANOVA, Tukey) Data represented as mean ± SD, n=3.

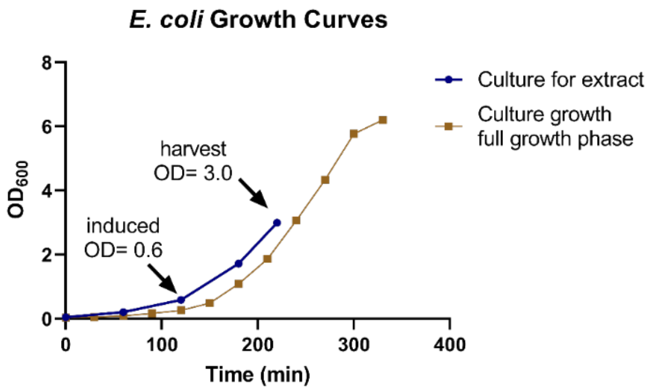


**Figure S2. Growth curves of BL21 DE3 Star *E. coli* strain used in this study.** The yellow line represents the growth curve of the BL21 when it is allowed to grow to a stationary phase to determine the harvesting OD point (mid-exponential phase, OD 3.0). The blue line represents the growth curve of the extract used, with the induction OD (0.6) and harvest OD (3.0) displayed.

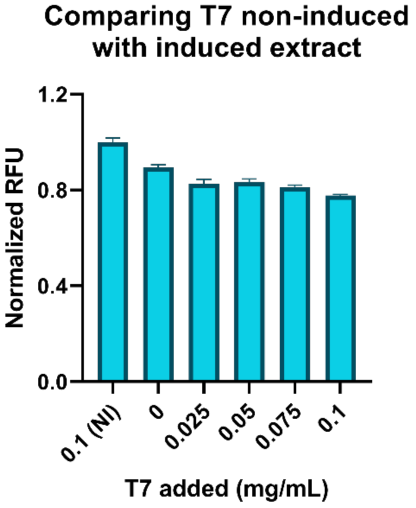


**Figure S3. T7 induced cell extract performance.** Non-induced (NI) T7 RNA polymerase (T7 RNAP) extract with added 0.1 mg/mL of purified T7 RNAP was compared to induced extract with different concentrations of additional purified T7 RNAP and with corresponding sfGFP fluorescence expression shown. Data represented as mean ± SD, n=3.

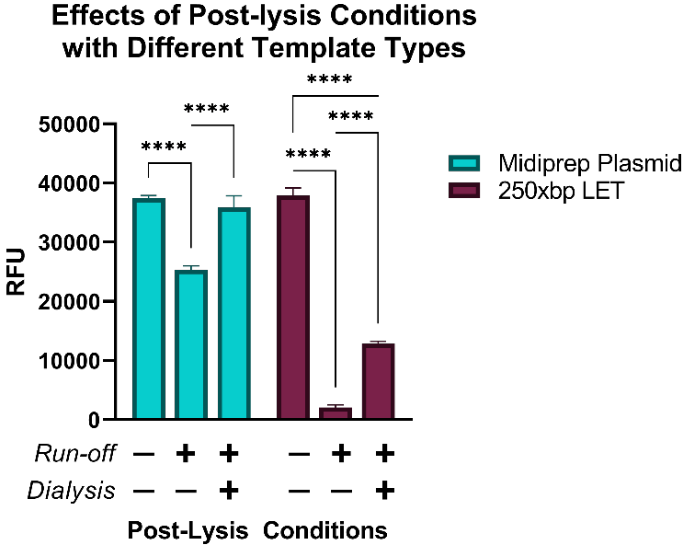


**Figure S4. Effects of post-lysis conditions on sfGFP protein production with varying DNA template types.** The BL21 lysate was subjected to dialysis and/or runoff reactions. Different DNA template types (midiprep blue, 250xbp LET red) were used in each post-lysis condition and protein production was measured. (***P<0.001, ****P<0.0001, two-way ANOVA, Tukey) Data represented as mean ± SD, n=3.

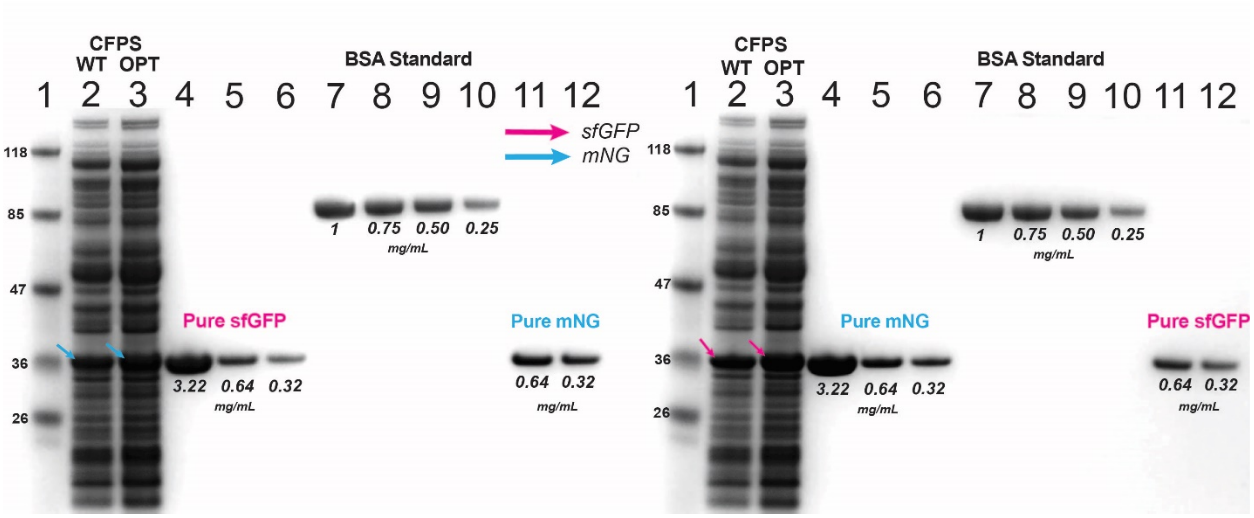


**Figure S5. mNG and sfGFP protein synthesis comparison between cell-free synthesized mNG and sfGFP with purified samples.** Proteins were analyzed using SDS-PAGE with Coomassie blue staining. L1: Protein standard ladder (kDa) L2&3: CFPS reaction total protein, comparing normal conditions (WT, L2) and optimal conditions (OPT, L3). mNG is pointed out with blue arrows on the left gel and sfGFP with pink arrows on the right gel. L4-L6: Strep-tag purified protein at different concentrations, mNG on the right gel, sfGFP on the left gel. L7-L10: BSA standard protein at different concentrations to visually compare to purified proteins. L11-L12: Purified proteins again at two different concentrations to compare the two on the same gel, mNG on the left gel, sfGFP on the right gel.

### Supplemental Methods

#### Buffers

| | Concentrations |
|---|---|
| **Buffer A** | 50 mM Tris, 14 mM magnesium glutamate, 60 mM potassium glutamate, pH 7.8 |
| **Buffer B** | 50 mM Tris, 14 mM magnesium glutamate, 60 mM potassium glutamate, 2 mM DTT, pH 7.8 |
| **Buffer C** | 5 mM Tris, 14 mM magnesium glutamate, 60 mM potassium glutamate, 1 mM DTT, pH 8.2 |

#### RBS calculator

The optimal RBS for every gene was found using the De Novo DNA web tool known as the RBS Calculator. The Design RBS algorithm was used by first setting the target translation initiation rate to "Maximize." This was done 5 separate times for each gene to find the sequence with the maximum translation rate. To make sure sfGFP and mNG had similar TIRs but still be at their maximum rate, the greatest rate out of the two was entered into the setting "target translation initiation rate" (in the same design algorithm tab) for the other, and the program was run 10 separate times until a sequence with the target translation initiation rate matched the first.

#### NUPACK

Secondary structures in the mRNA starting immediately after the promoter and 20 nucleotides into the open reading frame were analyzed using NUPACK. The settings were: Nucleic acid type-- RNA. Temperature-- 30.0 °C, number of strand species-- 1, maximum complex size-- 1. The sequence was then inserted into the text box and the "analyze" button was pressed. To edit the annotations on the output structure, "to utilities" was pressed.

#### Primers

Primer extension was used to add different RBS footprint sequences to sfGFP and mNG. Two primers were ordered for each gene that coded half complementary to the beginning of the gene and half of the new footprint and then half complementary to the new footprint already added and a half adding the rest footprint sequence. The primer conditions were set so that the one or two guanines or cytidines on the 5' end and one on the 3' end of the primer, melting temperature of the

3’ end around 47-40 °C, no runs of 4 or more of the same nucleotide, and the extension was around 15nt with the binding end also being around 15 nt.

Primer design for plasmid construction using Gibson Assembly also followed a set of rules. The primers were constructed with SnapGene by finding the junction point of the vector backbone and insert and following these guidelines as close as possible: a melting temperature of the binding region for PCR amplification around 55-60 °C, 15-20 nt on each side of the junction, GC content 35-65%, one or two guanines or a cytidines on the 5’ end and one on the 3’ end of the primer as anchors, and no runs of 4 or more of the same nucleotide.

**Detailed plasmid design**

PCR was carried out with the primers and templates with a cycle number of 30. The vector backbone was then treated with DpnI after PCR to degrade any mother plasmid. For extra caution, the vector backbone and the insert were gel extracted from a 2% agarose gel and purified with the Omega Bio-TEK, E.Z.N.A.® Gel Extraction Kit (V-spin) (D2500-01) and eluted in 60 °C nuclease-free ultrapure water (before elution, water was left to sit on the filter for 5 minutes).

The Gibson assembly mixture was made in-house containing 3.75% PEG-8000, 75 mM Tris-HCl (pH 7.5), 7.5 mM $MgCl_2$, 0.15 mM each dNTP, 7.5 mM DTT, 0.75 mM NAD, 0.004 U/µL T5 exonuclease, 0.025 U/µL Phusion polymerase, 4 U/µL Taq DNA Ligase, and nuclease-free water in the final reaction. The insert and vector were mixed with 2 µL of the Gibson Assembly Mix and incubated for 1 hour at 50 °C. 1 µL of Gibson Assembly mixture was transformed by electroporation into DH5α electrocompetent cells and then spread on kanamycin 50 µg/mL LB agar plates. The colonies were then grown for plasmid purification using the Omega Bio-TEK, E.Z.N.A.® Plasmid DNA Mini Kit I, (V-spin) (D6943-01). The concentration of the plasmids was measured using the BioTek Synergy HTX multi-mode reader and Take3 Micro-Volume Plate to determine a purity range of 1.8-1.9 with the 260/280 value and then the samples were stored in the -20 °C freezer. PCR and gel electrophoresis was performed to confirm the insert was there based on length. After, the plasmid inserts sequences were confirmed using Sanger Sequencing. For higher quality plasmid used in the cell-free protein synthesis reaction, Midiprep was done using the Qiagen kits.

**Cell-extract Dialysis and Runoff**

A run-off reaction was performed by incubating the clarified lysate at 37 °C with shaking at 200 rpm for 80 mins and then centrifugation was performed again to clarify the lysate (12,000 x g, 10 min, 4 °C). The supernatant was dialyzed using a 10K MWCO dialysis cassette in Buffer C at pH 8.2 for 3 hours at 4 °C. Then, the dialyzed lysate was centrifuged for a final time and the supernatant was collected and aliquoted out.

**Matrix effects CFPS experiment**

The cell-free protein synthesis reactions for the matrix effects experiments were built the same as the normal experiments, except a few things were added. 0.5 mg/mL of PVSA (Sigma-Aldrich, 278424) was added to the reaction to serve as an RNase inhibitor. The remaining volume (4.55 μL) was used for the sample providing the matrix effects. Tap water was taken from the faucet in the restroom. The pond water was taken from a small pond off-shot from the LSU lakes called "Campus Lake." The water was disturbed and then 0.5 L was taken with a bottle. The bottle was left undisturbed so that large particles would settle to the bottom, and then a 1 mL aliquot was taken. Whole milk was bought from the store and used. Hyclone Fetal bovine serum was filtered through a 0.4 μm filter and added to the reaction.

**Protein Purification**

pJL1-sfGFP and pJL1-mNG were transformed into BL21 DE3 Star. A colony was cultured at 37 °C, 250 rpm shaking overnight in 30 mL of LB media supplemented with kanamycin 50 μg/mL, 3. A 100 mL culture was started the following morning inoculated with 7 mL of the overnight culture and supplemented with kanamycin 50 μg/mL, 37 °C and 250 rpm shaking. The culture was grown until the OD reached 0.6 and then it was induced with IPTG so that the final concentration was 1 mM. The culture was then expressed for 6 hours. The culture was then harvested and frozen at -20 °C until the next day. The proteins were then purified by a strep-tag at the C-terminal of the protein using the Qiagen, Strep-Tactin Superflow Plus Cartridge. The manufacturer's protocol was followed for the remainder of the experiment. Once purified, protein concentration was determined using a BSA standard. A standard curve was also made for RFU given protein concentration using a series of dilutions. The dilutions were then added to a 96-well half-area black plate (Corning Incorporated, Corning, NY). Wells were filled with 5 μL of fluorescent protein dilutions and 45 μL of Milli-Q water. The plate was mixed in the plated reader orbitally at medium speed for 15 s and read at the height of 1.5 mm with a gain of 50. Another set of wells was filled and mixed the same and then measured at 48 gain. This way, a fluorescence standard curve could be made for both settings in the plate reader.
